# Timing and scope of genomic expansion within Annelida: evidence from homeoboxes in the genome of the earthworm *Eisenia fetida*

**DOI:** 10.1101/025130

**Authors:** Allison S Zwarycz, Carlos W Nossa, Nicholas H Putnam, Joseph F Ryan

**Affiliations:** Whitney Laboratory for Marine Bioscience, University of Florida, St. Augustine, FL 32080; Viterbo University, La Crosse, WI 54601; Department of Ecology and Evolutionary Biology, Rice University, Houston, TX 77005; Department of Biology, University of Florida, Gainesville, FL 32611

**Keywords:** Earthworm genome, Hox, Homeobox Evolution, Annelida, *Eisenia fetida*

## Abstract

Annelida represents a large and morphologically diverse group of bilaterian organisms. The recently published polychaete and leech genome sequences revealed an equally dynamic range of diversity at the genomic level. The availability of more annelid genomes will allow for the identification of evolutionary genomic events that helped shape the annelid lineage and better understand the diversity within the group. We sequenced and assembled the genome of the common earthworm, *Eisenia fetida*. As a first pass at understanding the diversity within the group, we classified 440 earthworm homeoboxes and compared them to those of the leech *Helobdella robusta* and the polychaete *Capitella teleta*. We inferred many gene expansions occurring in the lineage connecting the most recent common ancestor (MRCA) of *Capitella* and *Eisenia* to the *Eisenia*/*Helobdella* MRCA. Likewise, the lineage leading from the *Eisenia*/*Helobdella* MRCA to the leech *Helobdella robusta* has experienced substantial gains and losses. However, the lineage leading from *Eisenia*/*Helobdella* MRCA to *E. fetida* is characterized by extraordinary levels of homeobox gain. The evolutionary dynamics observed in the homeoboxes of these lineages are very likely to be generalizable to all genes. These genome expansions and losses have likely contributed to the remarkable biology exhibited in this group. These results provide a new perspective from which to understand the diversity within these lineages, show the utility of sub-draft genome assemblies for understanding genomic evolution, and provide a critical resource from which the biology of these animals can be studied. The genome data can be accessed through the *Eisenia fetida* Genome Portal: http://ryanlab.whitney.ufl.edu/genomes/Efet/

## Background

The remarkable variation among animal body plans can be attributed to historical innovations in the genomic components underlying animal development. The evolutionary dynamics of the homeobox superfamily in particular, have played an important role in the evolution of animal form (Lewis 1978; McGinnis, et al. 1984). Because of the well-known, highly conserved nature of the homeobox superfamily, we are able to distinguish genetic events that have taken place in the evolution of bilaterian species. Furthermore, analyzing the complete homeobox superfamily of a newly sequenced animal genome provides novel insight into the broader pattern of genomic evolution for that particular animal (Holland, et al. 2008; Martin and Holland 2014; Monteiro, et al. 2006; Paps, et al. 2015; Ryan, et al. 2006; Ryan, et al. 2010).

*Eisenia fetida* (also called red wigglers, compost worms, among other names) is a widespread non-burrowing earthworm that is known for its role in assessing terrestrial ecotoxicological levels (Spurgeon, et al. 1994). These worms are particularly well known for their ability to compost rotting material and are important for waste management and environmental monitoring. Their ecological importance was highlighted in Charles Darwin’s final book, *The Formation of Vegetable Mould through the Action of Worms* (Darwin 1892). *E. fetida* are highly amenable to laboratory manipulation and their success as a model system is clear from the more than 600 PubMed articles referencing *Eisenia fetida*. Despite the utility of *E. fetida*, very few molecular resources exist, apart from a limited set of expressed sequence tag (EST) data (Pirooznia, et al. 2007).

Like most other annelids, the basic body plan of *E. fetida* consists of a head followed by a segmented trunk and a tail. This simple body arrangement incorporates a tremendous amount of diversity including variation in number of segments, internal anatomy, as well as head and tail shapes. Homeobox transcription factors play a central patterning role during embryogenesis in most animals, and changes in the number, genomic arrangement, and regulation of these genes have been implicated in playing a major role in the diversification of animal body plans (Akam 1995). An important step in understanding the evolution of annelid body plan diversity is to understand the diversity of homeobox genes. There have been several studies of annelid homeobox genes (Andreeva, et al. 2001; Cho, et al. 2012; Dick and Buss 1994; Kulakova, et al. 2007), but most of these have considered only the HOXL subclass of genes and have concentrated on a single annelid species.

To date, complete genome sequences are available from two annelids: the marine polychaete *Capitella teleta* and the freshwater leech *Helobdella robusta* (Simakov, et al. 2013). The *C. teleta* genome is highly conserved in terms of genomic architecture (e.g., macrosynteny, intron-retention, gene retention, and gene duplication) when compared to other bilaterian genomes, while the *H. robusta* genome is considered relatively dynamic (Simakov, et al. 2013).

Notably, *E. fetida* shares a more recent common ancestor with *H. robusta* than either of them do with *C. teleta* (Erséus and Källersjö 2004; Purschke 2002; Weigert, et al. 2014) making it a useful model for understanding the timing of the dynamic genomic events that have occurred in the lineage leading to the leech. The presence, absence, and arrangement of Hox genes are prime examples of the hyper-dynamic nature of the *H. robusta* genome (Simakov, et al. 2013). Understanding the timing and frequency of these changes along the lineages leading to the leech and earthworm will shed light on how this shift has influenced the evolution of these two animals.

To this end, we have sequenced and assembled a draft-quality genome of the earthworm *Eisenia fetida*. Using this assembly, we are able to identify and phylogenetically classify the complete set of homeoboxes from *E. fetida*. Our analyses show that many of the gene duplication and loss events that are evident in the *H. robusta* genome predate the most recent common ancestor (MRCA) of *E. fetida* and *H. robusta*, and show that an extraordinary number of duplication events occurred in the earthworm lineage after it diverged from this ancestor.

## Methods

### Data Access

All genome sequencing data is available from the European Nucleotide Archive under the study accession: PRJEB10048 (http://www.ebi.ac.uk/ena/data/view/PRJEB10048). All alignments, trees, custom scripts, HMM models, and a detailed list of commands used in our analyses are available in our GitHub supplement: https://github.com/josephryan/RyanLab/tree/master/2015-Zwarycz_et_al

### Materials and Sequencing

Two adult, farm-raised *E. fetida* earthworms were crossed and produced 24 offspring. The digestive systems of the 26 worms were purged by being kept on moist paper clippings, out of dirt. The earthworms were washed with 70% ethanol prior to DNA extraction. DNA was extracted using Qiagen DNA Easy kit and Zymo Genomic DNA clean kit to further purify the DNA. Libraries were made with Nextera and sequenced on Illumina HiSeq2000 2 × 100PE. Sequencing was performed by the Beijing Genomics Institute (BGI).

### Error Correction and Adapter Trimming

Sequencing reads from each sample were concatenated and error correction was performed using the ErrorCorrectReads.pl program from Allpaths-LG version 44387 (Gnerre, et al. 2011). Besides read-cleaning, Allpaths-LG also estimates genome size based on k-mer spectrum. We used Cutadapt version 1.4.2 (Martin 2011) to remove adapter sequences from all error-corrected reads.

### Genome assembly

After adapter trimming and error correction, we created a total of ten genome assemblies using the following assemblers: SOAPdenovo version 2.04 (Luo, et al. 2012), ABySS version 3.81 (Simpson, et al. 2009), and Platanus version 1.2.1 (Kajitani, et al. 2014). Besides adjusting K-mer values, command-line parameters were mostly left as defaults.

### Assembly Evaluation

We evaluated each assembly using three primary criteria: (1) the number of *Eisenia fetida* expressed sequence tags (ESTs) that aligned to an assembly, (2) the number of 248 highly conserved eukaryotic genes (CEGs) identified with CEGMA version 2.4 (Parra, et al. 2007), and (3) the N50 statistic (Table 2). We used BLAT version 35x1 (Kent 2002) to align 4,329 *Eisenia fetida* ESTs available in GenBank (LIBEST_024375, LIBEST_026326, LIBEST_022256, and LIBEST_020813) and Isoblat version 0.3 (Ryan 2013) to gauge how well these ESTs mapped to each assembly.

We aligned the most ESTs using our ABySS assembly with K=63 (ABySS63 assembly), but this alignment produced the lowest CEGMA scores and had a very suboptimal N50. In contrast, we generated the highest CEGMA and N50 scores using our SOAPdenovo assembly with K=31 (SOAP31 assembly). The difference in number of mapped ESTs between the ABySS63 assembly and our SOAP31 assembly was negligible, whereas the differences in CEGMA scores and N50 values between these two assemblies were substantial. In addition, the size of the SOAP31 assembly was much closer to the Allpaths-LG prediction than the size of the ABySS63 assembly. Based on these results we chose the SOAP31 assembly for all downstream analyses.

### Homeodomain Dataset and Alignment

We ran the hmmsearch program from HMMer version 3.1b1 (Eddy 2011) on a translated version of our final assembly. For this search we used a custom homeobox hidden Markov model generated using hmmbuild on a FASTA file consisting of all homeodomains from HomeoDB (Zhong, et al. 2008) that were 60 amino acids in length (hd60.hmm in GitHub supplement). The resulting search produced an alignment (to the HMM) that we converted from STOCKHOLM to FASTA (using http://sequenceconversion.bugaco.com). We then removed all regions that did not align to the HMM (i.e., insertions) using remove_gaps_from_hmmsearch_results.pl (GitHub supplement). We repeated this process on filtered protein models of *C. teleta, H. robusta*, and *L. gigantea* that were downloaded from the Joint Genome Institute web site and from *C. gigas*, which was downloaded from GigaDB (Fang, et al. 2012). We labeled the sequences of *C. gigas* based on Paps et al. (2015).

For each species dataset, we used BLASTP (version 2.2.31+) and two custom Perl scripts to identify sequences that were missed in our initial HMMsearch runs. We first used our custom script hmmsearch_blast_combo.pl (GitHub supplement) to build a FASTA file with all of the sequences where a homeodomain was not recovered. We next ran a BLASTP with tabbed output and an e-value cutoff of 10 against all homeodomains from HomeoDB. We used our custom script parse_and_reblast_w_alignments.pl (GitHub supplement) to identify hits with E-Values below 0.001 and then to run a BLASTP search with default output on these searches. We extracted homeodomains by hand from BLAST alignments and then aligned them to the complete set of amphioxous homeodomains available from HomeoDB with MAFFT (v7.158b) and adjusted alignments by eye. We removed any insertions outside of the canonical 60 amino acid homeobox and then appended the non-amphioxous homeodomains to our grand set. In total, we added 26 *C. teleta*, 31 *H. robusta*, 17 *L. gigantea,* and 191 *E. fetida* homeodomains. The large number of additional *E. fetida* homeoboxes is due mostly to the lack of available protein models for this species. All of the homeodomains, both from the primary and secondary searches (1243 total) were included in our downstream analyses.

### Homeobox Phylogeny and Tree Generation

We used RAxML version 8.0.23 (Stamatakis 2006) to generate a maximum-likelihood (ML) tree from the aligned homeodomains of *E. fetida*, *C. teleta, H. robusta*, *L. gigantea,* and *C. gigas* (supplementary fig. S1). We pruned taxa from this tree with terminal branches longer than 2.3 using the custom script branch_lengths_filter.pl (GitHub supplement), which removed poorly predicted or extremely divergent homeodomains. This removed 29 sequences but left three sequences (Ct_214198, Ct_199162, Lg_132019) that appeared to be obviously false predictions (sequences available in GitHub supplement). These were manually removed. This pruning left us with 1,209 homeodomains, including 466 *Eisenia fetida*, 189 *Crassostrea gigas* (Mollusca), 155 *Lottia gigantea* (Mollusca), 271 *Helobdella robusta* (Annelida), and 178 *Capitella teleta* (Annelida) sequences.

We separated the 1,209 sequences into classes, as designated by HomeoDB (Zhong, et al. 2008) using the *C. gigas* class assignments as a guide. For each class-level dataset, we generated a maximum-likelihood tree with RAxML, corresponding bootstraps with the autoMRE stopping criteria in RAxML, and a Bayesian tree using MrBayes version 3.2.3 (Ronquist and Huelsenbeck 2003). Alignments and details for phylogenetic runs are in the GitHub supplement. For the Bayesian trees, the potential Scale Reduction Factor (PSRF) values produced by *sump* for both the sum of all branch lengths (TL), and the shape parameters of the gamma distribution of rate variation (alpha) were very close to one (the largest difference was for the TL in the Other analysis: PSRF=1.479514). According to the Mr. Bayes manual, PSRF values close to 1.0 suggest a good sample from the posterior probability distribution.

### Homeobox Classification and Naming

For each class, we computed a majority rule consensus tree using RAxML (with the -J STRICT option) from the ML and Bayesian trees. We used these consensus trees to classify each homeodomain at the family level in accordance with the family assigned to the *C. gigas* homeodomain in Paps et al. (2015). We examined homeodomains where the consensus trees were inconclusive by eye. In some cases, a homeodomain was excluded from a family clade in the consensus tree due to another homeodomain that was present in the clade in one of the trees but not the other. In these cases we classified these homeodomains as family members. In cases where two or more partial homeodomains were identified on the same scaffold in the same direction and not completely overlapping, we collapsed these into a single homeodomain. When more than one homeodomain from a single species was assigned to the same family, we used the family name, followed by a numerical label from their definition number. If we were unable to assign a homeodomain to a family, it was given the class/subclass name followed by “HD” and a number >20 (e.g., HOXLHD23). All homeodomain assignments are available in the supplemental material.

### Inferring Gene Duplications and Losses within Annelida

We used parsimony principles to infer the annelid evolutionary branch on which family-level gains and losses occurred. In this process, ancestral condition was estimated based on the number of homeodomains present in a *C. gigas* family, unless the particular *C. gigas* family was 0, in which case the *L. gigantea* number was used (in most cases these numbers were the same). We did not consider relationships within a family since support for intra-family relationships were mostly very low and would therefore require elaborate loss and gain scenarios. We classified each event (gains and losses) by determining the fewest number of events needed to explain the number of homeodomains identified in a particular family from the ancestral state. In cases where there were equally parsimonious explanations of the data, we chose the scenarios that maximized gains on external branches rather than internal branches. This biasing of events on terminal branches is justified based on the terminal branches of the three annelids all being over twice the length of the internal branch connecting the *Capitella*/Clitellata MRCA with the *Helobdella*/*Eisenia* MRCA (in Figure 1 of Weigert et al, 2014), suggesting that evolutionary events were more than twice as likely to occur on terminal branches.

**Fig. 1.**
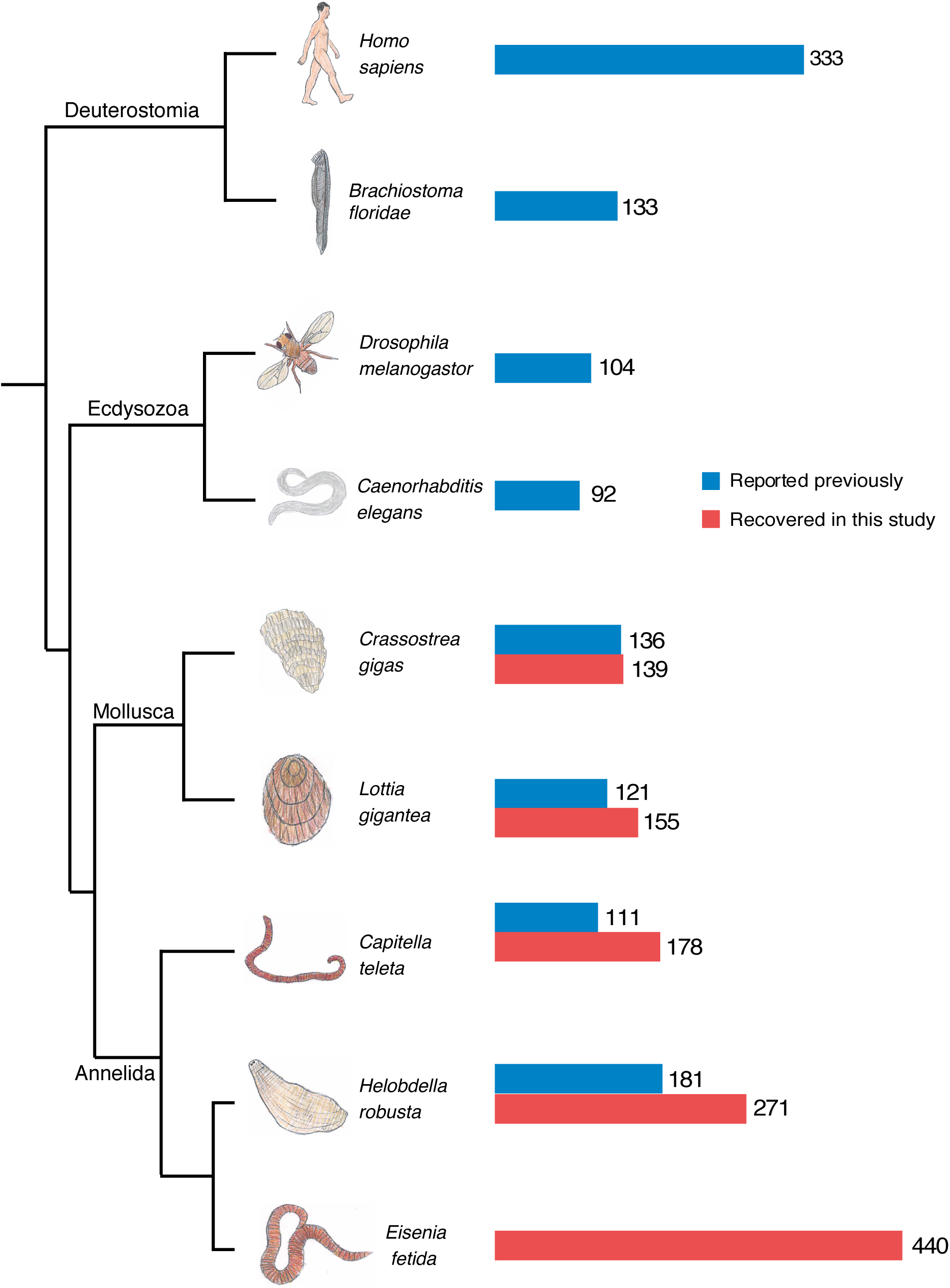
Phylogenetic relationship and total homeobox count of several bilaterian animals. The relationships of these taxa are based on multiple studies (Simakov, et al. 2013; Weigert, et al. 2014; Zhang, et al. 2012). The bar graph to the right of taxa labels shows the number of homeoboxes reported previously in blue, and the number of homeoboxes we identified in our analyses in red. *E. fetida* represents the highest reported total homeobox count among these animals.

## Results

We sequenced and assembled the genome of the earthworm *Eisenia fetida*. We identified 440 homeoboxes in this rough draft assembly. Furthermore, we retrieved 139 homeodomains from the oyster *C. gigas*, 155 from the limpet *L. gigantea*, 178 from the polychaete worm *C. teleta*, and 271 from the leech *H. robusta*. In all cases we were able to identify additional homeoboxes from each annelid and molluscan genome using our approach of combining HMM and BLAST approach (fig. 1). It should also be noted that a similar approach was used for *C. gigas* (Paps, et al. 2015), but that the others were discovered as part of whole-genome analyses, which were less targeted. We ran an extensive phylogenetic analysis using this comprehensive dataset as a means to understand the nature of the *E. fetida* genome, and some of the evolutionary dynamics that led to this genome. In the process, we classified the *E. fetida*, *C. teleta*, *H. robusta*, and *L. gigantea* homeoboxes based on careful designations applied to *C. gigas* in a recent comprehensive analysis of the homeoboxes of this animal (Paps, et al. 2015).

### Genome Assembly

We generated genomic reads from a mating pair of adult *E. fetida* and 24 offspring (100 bp paired-end reads on 300 bp inserts). After adapter trimming and error correction we generated 10 assemblies of these combined reads using three different assembly algorithms and a range of k-mer values. Our best assembly was 1.05 gigabases with an N50 of 1,850 bp (Table 1).

**Table 1.** *Eisenia fetida* assembly statistics.

| Program<br>(Parameters) | K-mer<br>Size | N50<br>(Bp) | Isoblat | CEGMA<br>(complete) | CEGMA<br>(partial) | Genome<br>Length (Gb) |
| --- | --- | --- | --- | --- | --- | --- |
| **SOAPdenovo**<br>(defaults, max<br>insert<br>size=300) | 31 | **1852** | 4103/4329 (95%) | **59** | **124** | 1.05 |
|  | 39 | 1086 | 4156/4329 (96%) | 53 | 110 | 1.28 |
|  | 45 | 781 | 4160/4329 (96%) | 46 | 105 | 1.47 |
|  | 55 | 375 | 4198/4329 (97%) | 39 | 91 | 2.07 |
|  | 63 | 422 | 4202/4329 (97%) | 36 | 93 | 2.13 |
| **ABYSS**<br>(defaults) | 31 | 61 | 4237/4329 (98%) | 30 | 74 | 2.22 |
|  | 45 | 94 | 4237/4329 (98%) | 18 | 67 | 2.41 |
|  | 63 | 165 | **4241/4329 (98%)** | 10 | 59 | 2.12 |
| **Platanus**<br>(defaults,<br>m=500) | 32 | 414 | 4108/4329 (95%) | 46 | 104 | 0.73 |
|  | 45 | 307 | 4119/4329 (95%) | 26 | 85 | 0.88 |

### Homeobox Phylogeny

From this assembly we isolated 466 homeoboxes. We also isolated homeoboxes from the following animals: *H. robusta* (271), *C. teleta* (178), *L. gigantea* (155), and *C. gigas* (139). In all cases we identified additional homeoboxes than had been identified in previous publications (fig. 1). From these data, we generated amino-acid alignments considering only the positions corresponding to the canonical 60 amino acid homeodomain.

### Class Designations

We ran a maximum-likelihood analysis using the complete set of homeodomains from the five animals in our study to classify each into one of the major classes or subclasses using the *C. Gigas* annotations as a guide. Figure 2 shows the distribution of homeoboxes according to class, with the most striking result being the large number of NKL, PRD, and LIM homeodomains in *E. fetida*. Similarly we found a major expansion of HOXL homeoboxes in both *E. fetida* and *H. robusta* (fig. 3). To classify these genes at the family level, we conducted both maximum likelihood and Bayesian phylogenetic analyses on each class with several smaller classes combined into a single group (other).

**Fig. 2.**
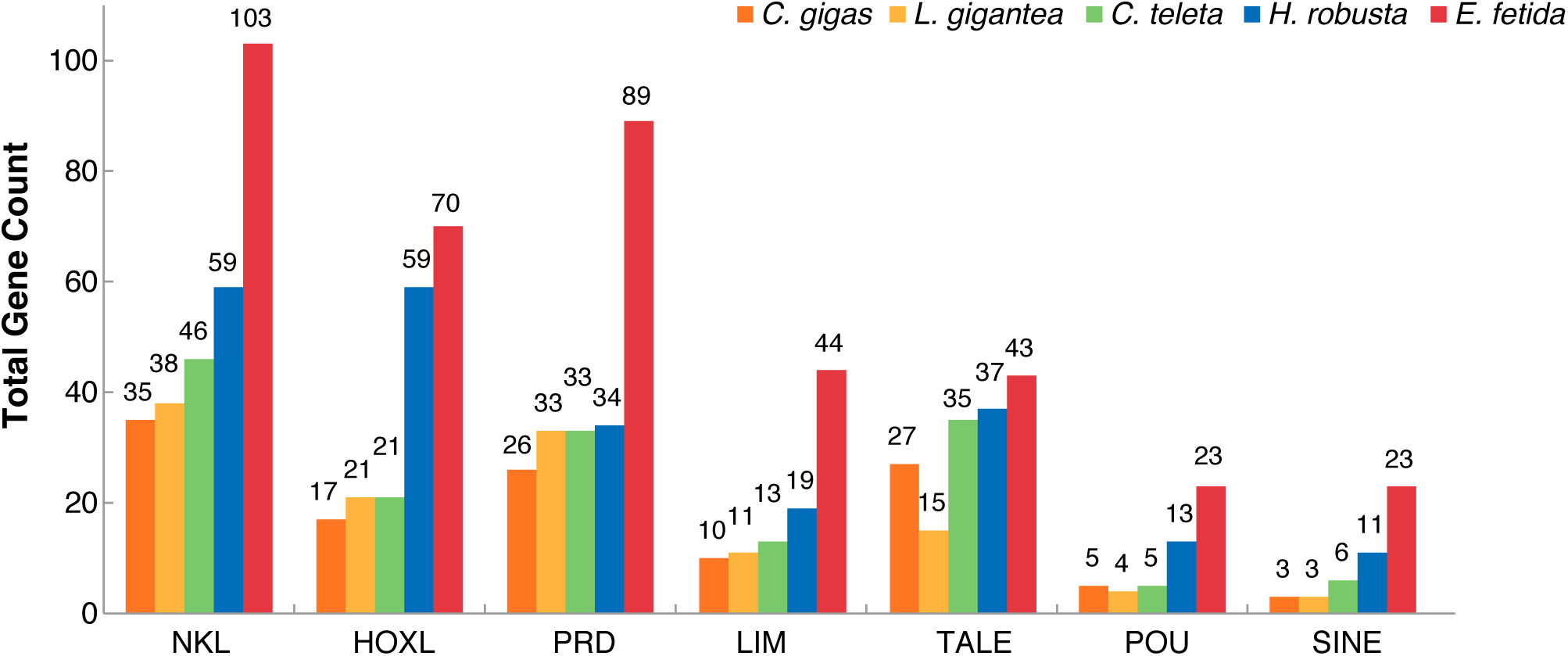
Total homeobox counts within classes/subclasses. Total homeobox counts for the largest homeobox classes (and subclasses) are shown for the molluscs *C. gigas* and *L. gigantea,* as well as the annelids *C. teleta*, *H. robusta*, and *E. fetida*. In all cases, *E. fetida* has the most homeoboxes, and H. *robusta* has the second-greatest number of homeoboxes relative all other animals in the comparison. Using a combination of searches with hidden Markov models (HMM) and BLAST, we added to the total homeoboxes that were found in previous studies.

**Fig. 3.**
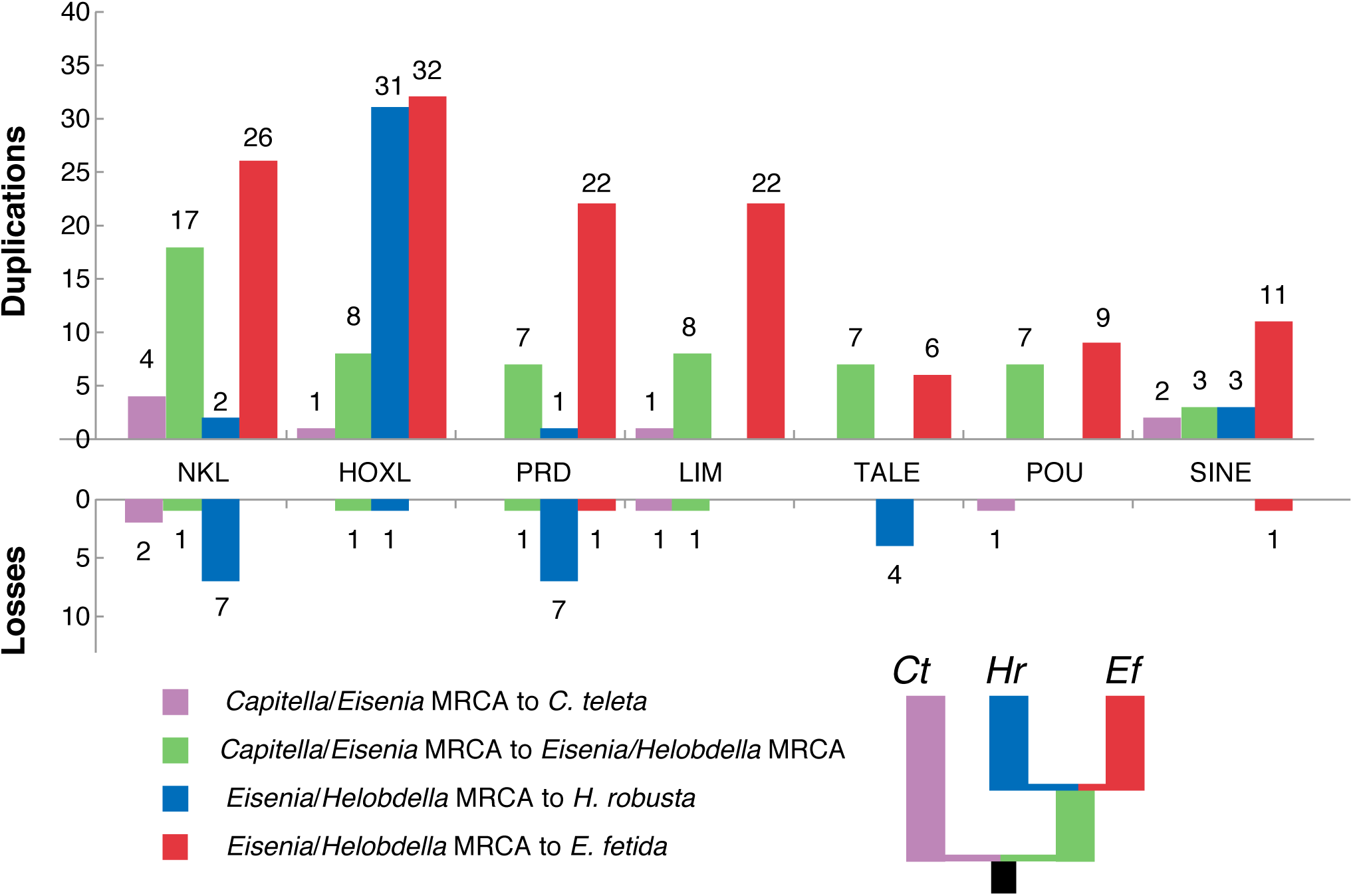
Inferred homeobox gains and losses in annelid lineages. Based on the distribution of homeobox genes on the phylogenetic tree in Fig. 1, we inferred gains and losses for four annelid lineages: (1) from the *Capitella*/*Eisenia* MRCA to *C. teleta*, (2) from the *Capitella*/*Eisenia* MRCA to the *Eisenia*/*Helobdella* MRCA, (3) from the *Eisenia*/*Helobdella* MRCA to *H. robusta*, and (4) from the *Eisenia*/*Helobdella* MRCA to *E. fetida*. The small tree next to the key shows lineage colors on the phylogeny.

### ANTP Class - HOXL Subclass

We identified 188 homeobox sequences belonging to the HOXL subclass of the ANTP class: 17 *C. gigas*, 21 *L. gigantea*, 21 *C. teleta*, 59 *H. robusta*, and 70 *E. fetida* (fig. 4; supplementary figs. S2-S4). The expansion of the HOXL complement in *H. robusta* is due mostly due to a clade of 28 homeoboxes that form a larger clade with CDX homeoboxes (in both Bayes and ML trees). If this placement is true, it appears that the CDX gene duplicated 28 times in the lineage leading to *H. robusta* after diverging from the *E. fetida* lineage. This is an unprecedented degree of duplication in a Hox/ParaHox gene family. Besides the CDX duplications, we inferred eight duplication events in six HOXL families that occurred along the lineage leading to the *H. robusta*/*E. fetida* MRCA after the split from *C. teleta*. We also inferred 32 duplication events in 11 HOXL families in the *E. fetida* lineage after the split from *H. robusta* (figs. 4-5). We identify a loss of the Pb Hox gene in the lineage leading to the MRCA of *H. robusta* and *E. fetida*, and show that the Hox3 homeobox was lost in the lineage leading to *H. robusta* after diverging from *E. fetida*. Incidentally, Hox3 is reported to be present in *H. robusta* in Simakov et al. (2012), but we could not identify it in our analyses.

**Fig. 4.**
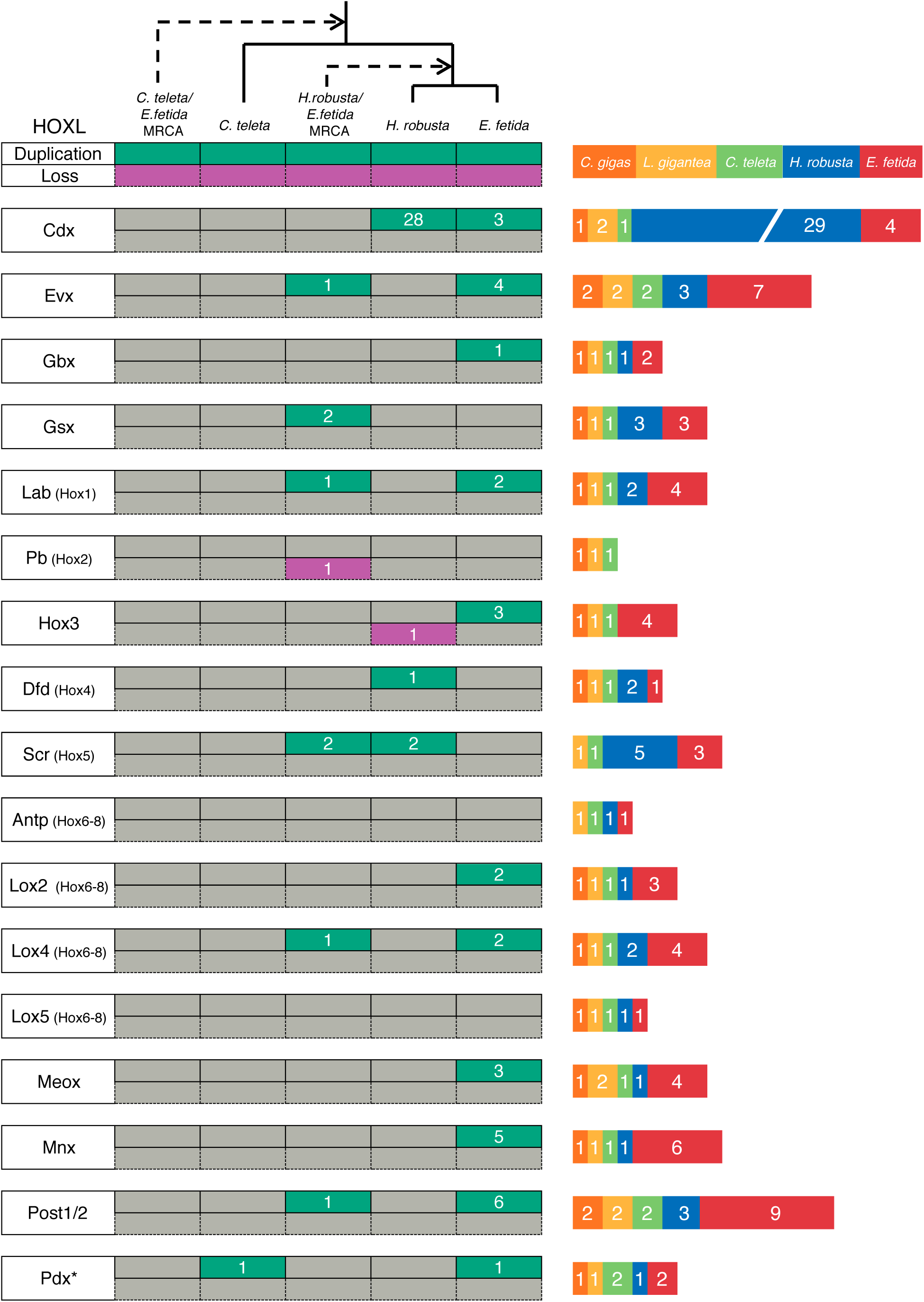
HOXL subclass (ANTP) evolutionary events and total homeobox count. Each green box on the left side of the figure represents one or more inferred duplication events, while each purple box represents one or more inferred losses. The number inside each green or purple box indicates the number of genes inferred to be gained or lost. An asterisk indicates more than one possible transition leading to the final homeobox count. The stacked bar graph represents the number of homeboxes in each family for each species. Zero values do not appear in the graph. The dashed lines indicate ancestral nodes.

### ANTP Class - NKL Subclass

There was a major expansion of NKL homeoboxes (26 in 13 families) in the lineage leading to *E. fetida* after the *Eisenia/Helobdella* MRCA. We also inferred 17 gains of NKL homeoboxes in the lineage leading to the *Eisenia/Helobdella* MRCA and seven losses in the lineage leading to *H. robusta* from this ancestor. Lastly, we inferred the loss of four NKL-class homeoboxes in the lineage leading to the Capitellidae/Clitellata ancestor, which includes the Vax, Nk4, Msx, and Hlx families (supplementary figs. S5-S8).

### PRD Class

As in the ANTP class, we deduced in the PRD class an extraordinary number of homeobox gene duplications (22 in nine families) to have occurred in the lineage leading to *E. fetida* after diverging from the *Eisenia/Helobdella* MRCA. In addition, we inferred seven gains in the *Eisenia/Helobdella* ancestor in six different families (supplementary fig. S9-S12). We inferred seven losses in the lineage leading to *H. robusta* after the *Eisenia/Helobdella* MRCA and also the loss of the Pax4/6 family in the lineage leading to the Capitellidae/Clitellata ancestor.

### LIM Class

We inferred that the number of LIM class homeobox genes more than doubled in the lineage leading to *E. fetida* after the *Eisenia/Helobdella* MRCA (22 duplications occurring in seven of the nine families). There were also six duplications in the Islet family that occurred in the lineage leading to the *Eisenia/Helobdella* MRCA after the split from *Capitella* (supplementary figs. S13-16).

### Other Classes

We have highlighted the largest homeobox classes, all of which experienced major expansions in the *E. fetida* lineage. Most of the other classes also experienced expansions in the *E. fetida* lineage, especially the SINE class where we inferred 11 duplications in the lineage leading to *E. fetida* after the *Eisenia/Helobdella* MRCA (supplementary figs. S17-S32).

## Discussion

We have sequenced and assembled the genome of the earthworm *Eisenia fetida*. We have also annotated 440 *E. fetida* homeoboxes in an effort to understand the evolutionary dynamics that shaped its genome. We have inferred the timing of an extraordinary number of evolutionary genomic events in the form of homeobox gene duplications and losses that have occurred in the lineage leading to *E. fetida*.

### Rates of Gene Gain and Gene Loss

Consistent with previous studies (Simakov, et al. 2013), we find that numerous homeobox gains and losses occurred in the lineage leading from the Capitellidae/Clitellata ancestor to the lineage leading to *H. robusta*. Using data from the earthworm genome, we show that many of these events occurred prior to the most recent common ancestor of *H. robusta* and *E. fetida*, and that while the *H. robusta* lineage continued to experience additional homeobox gains and losses after the split from this ancestor, the gene duplication process was accelerated in the lineage leading to *E. fetida*. Based on the pattern that we observe in the homeobox superfamily of genes, we propose that dynamics of genomic gain and loss rates is generalizable across each of these annelid lineages.

### E. fetida Hox Genes

We recovered 30 Hox genes from the *E. fetida* genome (Fig. 5). It will be interesting to see the extent of clustering in the *E. fetida* Hox genes once there is higher genomic resolution. This resolution will provide some insight into the nature of gene expansion (i.e., individual duplications vs. large segmental duplications) in these lineages. For example, if these 30 Hox genes are situated in multiple Hox clusters, it would suggest that gene expansion in the *E. fetida* lineage was due to large segmental duplications and possibly whole-genome duplication(s).

**Fig. 5.**
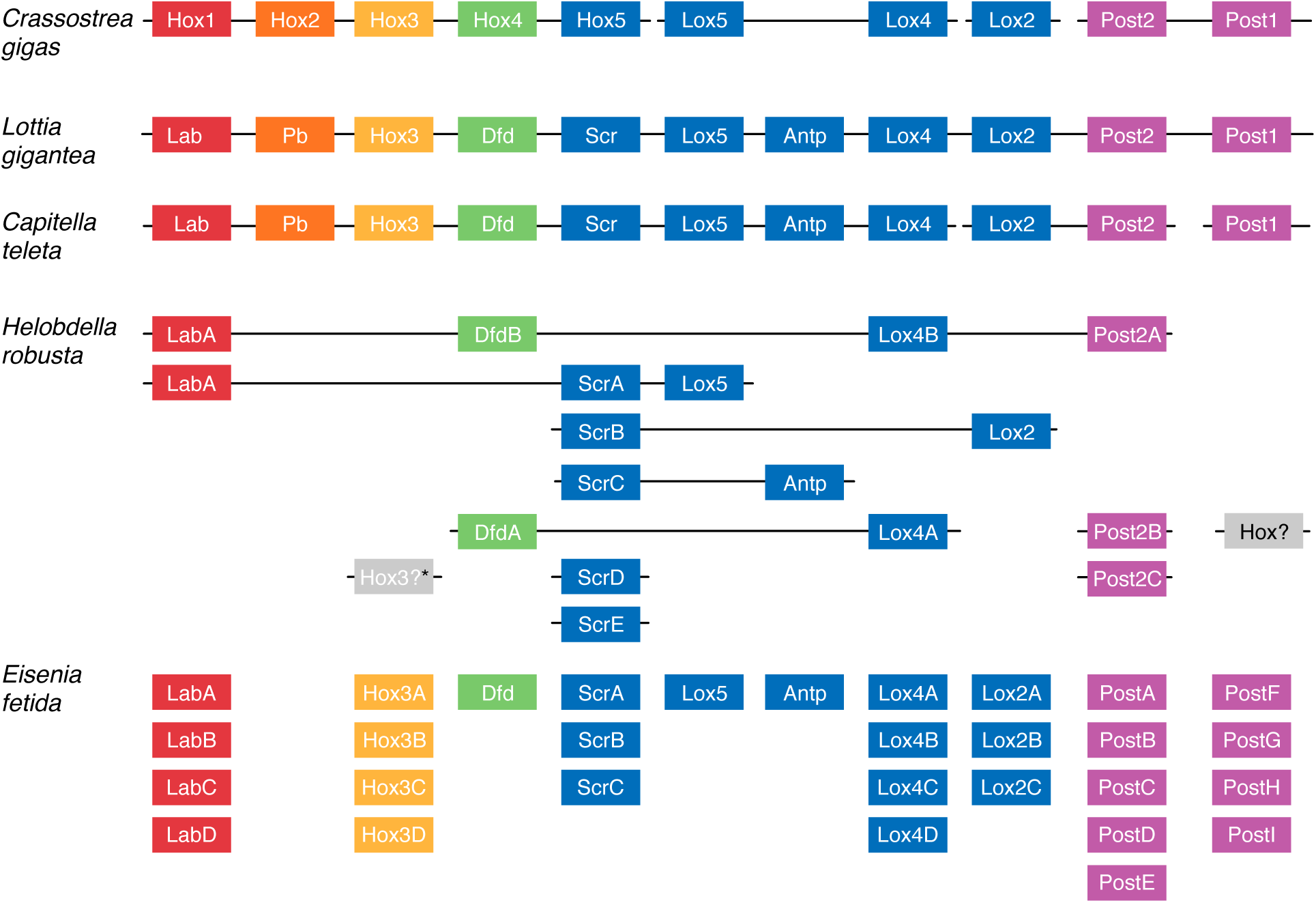
Hox clustering and count across two molluscs and three annelids. The *C. gigas* Hox cluster was obtained from Zhang et al. 2012. The *L. gigantea, H. robusta* and *C. teleta* clusters were obtained from Simakov et al. 2012. *C. gigas*, *L. gigantea,* and *C. teleta* maintain a single copy of each Hox homeobox while both *H. robusta* and *E. fetida* show multiple duplications in several families. The Post families experienced the largest number of duplications. However, we were unable to distinguish between the Post1 and Post2 in *E. fetida*. *Although a *H. robusta* Hox3 gene was identified in Simakov et al. (2012), we were unable to identify it in our analyses.

### Utility of Sub-Draft-Level Genomes

By using a sub-draft level genome, we were able to identify an unprecedented number of homeoboxes in an invertebrate animal and in the process, better understand how annelid genomes have evolved. Assembling large (> 1 gigabase) genome sequences to a high quality level requires the use of multiple technologies (e.g Illumina paired-end sequencing + Fosmids, BACs, PacBio, mate pairs, etc.), a large team and is expensive and time consuming. We were able to generate a sub-draft level genome assembly using only Illumina technology and a small team of researchers in a reasonable amount of time. It is clear from these analyses that genome sequences of this quality can be useful for understanding broad principles in genome evolution, and we have shown that this approach is able to provide a better understanding of animal genome dynamics within a localized clade of animals. By scaling this approach, it is possible to gain a broad evolutionary perspective of how genomes have evolved across animals, and possibly reveal general evolutionary principles.

### Implications of Gene Duplications on E. fetida Embryogenesis

Expanding taxon sampling will be critical for determining more precise timing for the described genomic expansion events. Genome sequencing of additional ingroup taxa (e.g., annelids from within Terebelliformia, Arenicolidae, Opheliidae, and Echiura) is a necessary step towards better resolution of the timing of events. This will in turn be critical for correlating events with the origin of synapomorphies in various clades of Sedentaria (a clade of annelids that include *Capitella*, *Helobdella*, and *Eisenia*) (Weigert, et al. 2014).

Annelids (and many other spiralians) have a highly stereotyped cleavage program called spiral cleaveage (Shankland and Seaver 2000). Earthworms have a spiral cleavage program that is a significant departure from other annelids (Anderson 2013). Homeobox genes are critical throughout embryogenesis including early development (Carroll, et al. 2013). This huge expansion in these important developmental genes represents a prime target for research into understanding the changes in early earthworm development. The *E. fetida* genome sequence provides an indispensible tool from which the genetic causes of this developmental diversity can be investigated.

## Acknowledgements

We thank Marta Chiodin, Daniel Sasson, and Elaine Seaver for helpful comments on previous versions of this manuscript. ASZ received support from the National Science Foundation Research Experience For Undergraduates (REU) Program (DBI-1156528).

## Supplementary Figure Legends

**Supplementary Fig. 1. Maximum-likelihood tree of all homeodomains in this study.** Cg = *C.gigas*, Lg = *L. gigantea*, Ct = *C. teleta*, Hr = *H. robusta*, Ef = *E. fetida*.

**Supplementary Fig. 2. Maximum-likelihood tree of HOXL homeodomains.** Numbers at the nodes are bootstrap values based on 650 bootstraps. Cg = *C.gigas*, Lg = *L. gigantea*, Ct = *C. teleta*, Hr = *H. robusta*, Ef = *E. fetida*.

**Supplementary Fig. 3. Bayesian tree of HOXL homeodomains.** Numbers at the nodes are posterior probabilities represented as percentages. Cg = *C.gigas*, Lg = *L. gigantea*, Ct = *C. teleta*, Hr = *H. robusta*, Ef = *E. fetida*.

**Supplementary Fig. 4. Maximum-likelihood/Bayesian Consensus tree of HOXL homeodomains.** Cg = *C.gigas*, Lg = *L. gigantea*, Ct = *C. teleta*, Hr = *H. robusta*, Ef = *E. fetida*.

**Supplementary Fig. 5. NKL subclass (ANTP) evolutionary events and total homeobox count.** Each green box on the left side of the figure represents one or more inferred duplication events, while each purple box represents one or more inferred losses. The number inside each green or purple box indicates the number of genes inferred to be gained or lost. An asterisk indicates more than one possible transition leading to the final homeobox count. The stacked bar graph represents the number of homeboxes in each family for each species. Zero values do not appear in the graph.

**Supplementary Fig. 6. Maximum-likelihood tree of NKL homeodomains.** Numbers at the nodes are bootstrap values based on 600 bootstraps. Cg = *C.gigas*, Lg = *L. gigantea*, Ct = *C. teleta*, Hr = *H. robusta*, Ef = *E. fetida*.

**Supplementary Fig. 7. Bayesian tree of NKL homeodomains.** Numbers at the nodes are posterior probabilities represented as percentages. Cg = *C.gigas*, Lg = *L. gigantea*, Ct = *C. teleta*, Hr = *H. robusta*, Ef = *E. fetida*.

**Supplementary Fig. 8. Maximum-likelihood/Bayesian Consensus tree of NKL homeodomains.** Cg = *C.gigas*, Lg = *L. gigantea*, Ct = *C. teleta*, Hr = *H. robusta*, Ef = *E. fetida*.

**Supplementary Fig. 9. PRD class evolutionary events and total homeobox count.** Each green box on the left side of the figure represents one or more inferred duplication events, while each purple box represents one or more inferred losses. The number inside each green or purple box indicates the number of genes inferred to be gained or lost. An asterisk indicates more than one possible transition leading to the final homeobox count. The stacked bar graph represents the number of homeboxes in each family for each species. Zero values do not appear in the graph.

**Supplementary Fig. 10. Maximum-likelihood tree of PRD class homeodomains.** Numbers at the nodes are bootstrap values based on 600 bootstraps. Cg = *C.gigas*, Lg = *L. gigantea*, Ct = *C. teleta*, Hr = *H. robusta*, Ef = *E. fetida*.

**Supplementary Fig. 11. Bayesian tree of PRD class homeodomains.** Numbers at the nodes are posterior probabilities represented as percentages. Cg = *C.gigas*, Lg = *L. gigantea*, Ct = *C. teleta*, Hr = *H. robusta*, Ef = *E. fetida*.

**Supplementary Fig. 12. Maximum-likelihood/Bayesian Consensus tree of PRD class homeodomains.** Cg = *C.gigas*, Lg = *L. gigantea*, Ct = *C. teleta*, Hr = *H. robusta*, Ef = *E. fetida*.

**Supplementary Fig. 13. LIM class evolutionary events and total homeobox count.** Each green box on the left side of the figure represents one or more inferred duplication events, while each purple box represents one or more inferred losses. The number inside each green or purple box indicates the number of genes inferred to be gained or lost. An asterisk indicates more than one possible transition leading to the final homeobox count. The stacked bar graph represents the number of homeboxes in each family for each species. Zero values do not appear in the graph.

**Supplementary Fig. 14. Maximum-likelihood tree of LIM class homeodomains.** Numbers at the nodes are bootstrap values based on 900 bootstraps. Cg = *C.gigas*, Lg = *L. gigantea*, Ct = *C. teleta*, Hr = *H. robusta*, Ef = *E. fetida*.

**Supplementary Fig. 15. Bayesian tree of LIM class homeodomains.** Numbers at the nodes are posterior probabilities represented as percentages. Cg = *C.gigas*, Lg = *L. gigantea*, Ct = *C. teleta*, Hr = *H. robusta*, Ef = *E. fetida*.

**Supplementary Fig. 16. Maximum-likelihood/Bayesian Consensus tree of LIM class homeodomains.** Cg = *C.gigas*, Lg = *L. gigantea*, Ct = *C. teleta*, Hr = *H. robusta*, Ef = *E. fetida*.

**Supplementary Fig. 17. SINE class evolutionary events and total homeobox count.** Each green box on the left side of the figure represents one or more inferred duplication events, while each purple box represents one or more inferred losses. The number inside each green or purple box indicates the number of genes inferred to be gained or lost. An asterisk indicates more than one possible transition leading to the final homeobox count. The stacked bar graph represents the number of homeboxes in each family for each species. Zero values do not appear in the graph.

**Supplementary Fig. 18. Maximum-likelihood tree of SINE class homeodomains.** Numbers at the nodes are bootstrap values based on 1000 bootstraps. Cg = *C.gigas*, Lg = *L. gigantea*, Ct = *C. teleta*, Hr = *H. robusta*, Ef = *E. fetida*.

**Supplementary Fig. 19. Bayesian tree of SINE class homeodomains.** Numbers at the nodes are posterior probabilities represented as percentages. Cg = *C.gigas*, Lg = *L. gigantea*, Ct = *C. teleta*, Hr = *H. robusta*, Ef = *E. fetida*.

**Supplementary Fig. 20. Maximum-likelihood/Bayesian Consensus tree of SINE class homeodomains.** Cg = *C.gigas*, Lg = *L. gigantea*, Ct = *C. teleta*, Hr = *H. robusta*, Ef = *E. fetida*.

**Supplementary Fig. 21. POU class evolutionary events and total homeobox count.** Each green box on the left side of the figure represents one or more inferred duplication events, while each purple box represents one or more inferred losses. The number inside each green or purple box indicates the number of genes inferred to be gained or lost. An asterisk indicates more than one possible transition leading to the final homeobox count. The stacked bar graph represents the number of homeboxes in each family for each species. Zero values do not appear in the graph.

**Supplementary Fig. 22. Maximum-likelihood tree of POU class homeodomains.** Numbers at the nodes are bootstrap values based on 1000 bootstraps. Cg = *C.gigas*, Lg = *L. gigantea*, Ct = *C. teleta*, Hr = *H. robusta*, Ef = *E. fetida*.

**Supplementary Fig. 23. Bayesian tree of POU class homeodomains.** Numbers at the nodes are posterior probabilities represented as percentages. Cg = *C.gigas*, Lg = *L. gigantea*, Ct = *C. teleta*, Hr = *H. robusta*, Ef = *E. fetida*.

**Supplementary Fig. 24. Maximum-likelihood/Bayesian Consensus tree of POU class homeodomains.** Cg = *C.gigas*, Lg = *L. gigantea*, Ct = *C. teleta*, Hr = *H. robusta*, Ef = *E. fetida*.

**Supplementary Fig. 25. TALE class evolutionary events and total homeobox count.** Each green box on the left side of the figure represents one or more inferred duplication events, while each purple box represents one or more inferred losses. The number inside each green or purple box indicates the number of genes inferred to be gained or lost. An asterisk indicates more than one possible transition leading to the final homeobox count. The stacked bar graph represents the number of homeboxes in each family for each species. Zero values do not appear in the graph.

**Supplementary Fig. 26. Maximum-likelihood tree of TALE class homeodomains.** Numbers at the nodes are bootstrap values based on 500 bootstraps. Cg = *C.gigas*, Lg = *L. gigantea*, Ct =*C. teleta*, Hr = *H. robusta*, Ef = *E. fetida*.

**Supplementary Fig. 27. Bayesian tree of TALE class homeodomains.** Numbers at the nodes are posterior probabilities represented as percentages. Cg = *C.gigas*, Lg = *L. gigantea*, Ct = *C. teleta*, Hr = *H. robusta*, Ef = *E. fetida*.

**Supplementary Fig. 28. Maximum-likelihood/Bayesian Consensus tree of TALE class homeodomains.** Cg = *C.gigas*, Lg = *L. gigantea*, Ct = *C. teleta*, Hr = *H. robusta*, Ef = *E. fetida*.

**Supplementary Fig. 29. Other classes evolutionary events and total homeobox count.** This group includes multiple classes that are not necessarily monophyletic. The classes represented include ZF, CERS, PROS, and CUT. Each green box on the left side of the figure represents one or more inferred duplication events, while each purple box represents one or more inferred losses. The number inside each green or purple box indicates the number of genes inferred to be gained or lost. An asterisk indicates more than one possible transition leading to the final homeobox count. The stacked bar graph represents the number of homeboxes in each family for each species. Zero values do not appear in the graph.

**Supplementary Fig. 30. Maximum-likelihood tree of all other homeodomains.** Numbers at the nodes are bootstrap values based on 500 bootstraps. Cg = *C.gigas*, Lg = *L. gigantea*, Ct = *C. teleta*, Hr = *H. robusta*, Ef = *E. fetida*.

**Supplementary Fig. 31. Bayesian tree of all other homeodomains.** Numbers at the nodes are posterior probabilities represented as percentages. Cg = *C.gigas*, Lg = *L. gigantea*, Ct = *C. teleta*, Hr = *H. robusta*, Ef = *E. fetida*.

**Supplementary Fig. 32. Maximum-likelihood/Bayesian Consensus tree of all other homeodomains.** Cg = *C.gigas*, Lg = *L. gigantea*, Ct = *C. teleta*, Hr = *H. robusta*, Ef = *E. fetida*.

